# *SCARECROW* gene function is required for photosynthetic development in maize

**DOI:** 10.1101/2020.06.28.176305

**Authors:** Thomas E. Hughes, Jane A. Langdale

## Abstract

C_4_ photosynthesis in grasses relies on a specialized leaf anatomy. In maize, this ‘Kranz’ leaf anatomy is patterned in part by the duplicated *SCARECROW (SCR)* genes *ZmSCR1* and *ZmSCR1h*. Here we show that in addition to patterning defects, chlorophyll content and levels of transcripts encoding Golden2-like regulators of chloroplast development are significantly lower in *Zmscr1;Zmscr1h* mutants than in wild-type. These perturbations are not associated with changes in chloroplast number, size or ultrastructure. However, the maximum rates of carboxylation by ribulose bisphosphate carboxylase/oxygenase (RuBisCO, V_cmax_) and phospho*enol*pyruvate carboxylase (PEPC, V_pmax_) are both reduced, leading to perturbed plant growth. The CO_2_ compensation point and ^13^C‰ *of Zmscr1;Zmscr1h* plants are both normal, indicating that a canonical C_4_ cycle is operating, albeit at reduced overall capacity. Taken together, our results reveal that the maize *SCR* genes, either directly or indirectly, play a role in photosynthetic development.

**Significance statement:** SCARECROW (SCR) is one of the best studied plant developmental regulators, however, its role in downstream plant physiology is less well-understood. Here, we have demonstrated that SCR is required to establish and/or maintain photosynthetic capacity in maize leaves.

## Introduction

The C_4_ photosynthetic pathway is nearly always underpinned by characteristic Kranz anatomy, whereby vascular bundles are closely spaced and are often separated by only two mesophyll (M) cells (reviewed in Sedelnikova et al., 2018). This cellular arrangement enables function of the C_4_ cycle, in which CO_2_ is initially fixed in the outer M cells by phospho*enol*pyruvate carboxylase (PEPC) into the 4-carbon compound oxaloacetate. Although there are at least three distinct C_4_ enzymatic cycles (Furbank, 2011), all involve a 4-carbon derivative of oxaloacetate being shuttled to the inner bundle-sheath (BS) cells to be decarboxylated. The CO_2_ released in the BS cells is refixed by ribulose bisphosphate carboxylase/oxygenase (RuBisCO) in the Calvin-Benson (C_3_) cycle. Importantly, refixation occurs in a cellular environment where local levels of CO_2_ are high enough to suppress the competing oxygenation reaction and thus to prevent the energetically wasteful process of photorespiration. As compared to ancestral C_3_ species, the carbon concentrating mechanism that operates in C_4_ species can lead to higher crop yields, as well as to improvements in water and nitrogen use efficiency (Brown, 1999; Sheehy et al., 2007).

Developmental regulators of Kranz anatomy have proved elusive; however, it was recently reported that the duplicated maize *SCARECROW (SCR)* genes *ZmSCR1* and *ZmSCR1h* redundantly regulate Kranz patterning (Hughes et al., 2019). In *Zmscr1;Zmscr1h* mutants, the majority of vascular bundles are separated by only one rather than two mesophyll cells, and there are additional patterning perturbations in both leaves and roots (Hughes et al., 2019). Roles for *SCR* genes in patterning processes are well established, particularly in *Arabidopsis thaliana* (hereafter referred to as Arabidopsis), in which *AtSCR* (the single copy ortholog of *ZmSCR1* and *ZmSCR1h*) regulates the development of both the endodermis in the root and the BS in the leaf (Cui et al., 2014; Di Laurenzio et al., 1996; Wysocka-Diller et al., 2000). In addition to patterning defects, both *Atscr* and *Zmscr1;Zmscr1h* mutants are smaller than wild-type (Dhondt et al., 2010; Hughes et al., 2019), with reduced plant size In *Atscr* mutants associated with reduced cell proliferation rather than impaired root function (Dhondt et al., 2010). The effect of SCR on other downstream developmental and physiological processes, such as chloroplast development, photosynthesis and growth, is less well understood.

To further investigate the role of SCR in shoot growth and development, we undertook a physiological characterisation of *Zmscr1;Zmscr1h* mutant leaves. We show here that chlorophyll levels are reduced in *Zmscr1;Zmscr1h* double mutants, but not in either of the corresponding single mutants. Although transcript levels of the chloroplast developmental regulators *ZmGLK1* and *ZmG2* are reduced in *Zmscr1;Zmscr1h* leaves, there are no apparent perturbations to chloroplast development. The CO_2_ compensation point and ^13^C ‰ *of Zmscr1;Zmscr1h* leaves are also normal, suggesting that the C_4_ photosynthetic cycle is operational, albeit with reduced PEPC fixation capacity and reduced maximum photosynthetic rate. Collectively, these results imply a role for ZmSCR1 and ZmSCR1h (either directly or indirectly) in the establishment and/or maintenance of photosynthetic capacity in the maize leaf.

## Results

### Chlorophyll levels are reduced in *Zmscr1;Zmscr1h* mutants

*Zmscr1;Zmscr1h* plants appear paler green than either wild-type or single mutants. To quantify this difference, the chlorophyll content in multiple individuals from each genotype was measured (Figure 1). In general, overall chlorophyll levels in WT (*m2m1*) (a segregating wild-type line from the *m2m1* F2 population) and single mutant leaves were similar, although levels were slightly lower in the *Zmscr1h-m2* line (Fig 1A). In contrast, both *Zmscr1-m2;Zmscr1h-m1* and *Zmscr1-m2;Zmscr1h-m2* mutants (hereafter referred to as the *m2m1* and *m2m2* lines for simplicity) had significantly lower levels of total chlorophyll than WT (Fig 1A). When analysed individually, both chlorophyll a and chlorophyll b levels were reduced by a similar magnitude in *Zmscr1;Zmscr1h* mutants (Fig 1B,C). The pale green phenotype of *Zmscr1;Zmscr1h* double mutants is thus caused by a 52% (*m2m1*) or 42% (*m2m2*) reduction in total chlorophyll levels, a defect that is complemented by activity of either ZmSCR1 or ZmSCR1h in single mutants. All subsequent analyses therefore focussed on comparisons between wild-type and double mutant plants.

**Figure 1.**
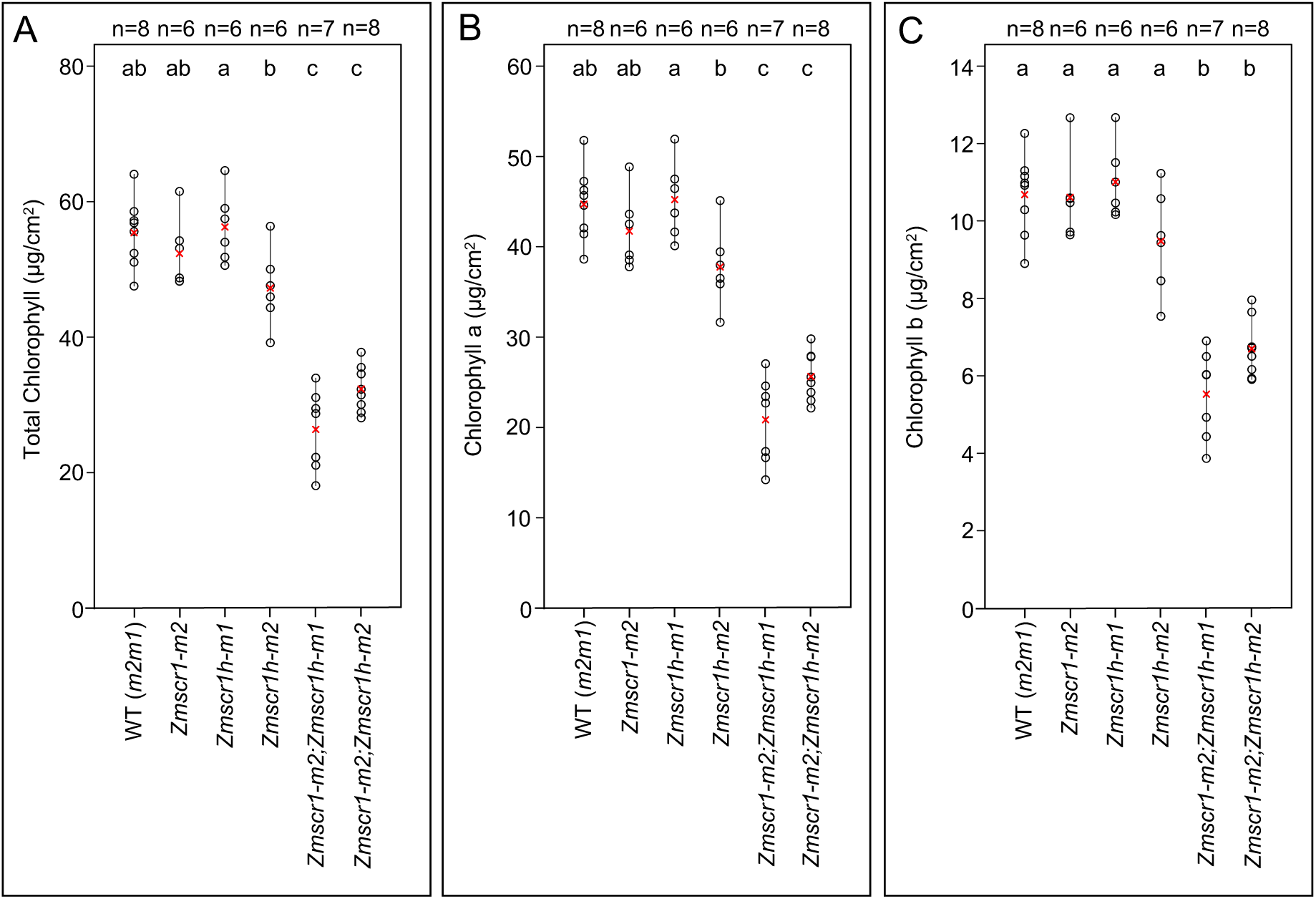
Chlorophyll levels are reduced in *Zmscr1;Zmscr1h* mutants. Total chlorophyll (A), chlorophyll a (B) and chlorophyll b (C) expressed on a leaf area basis. Biological replicates (n=) are shown above each genotype. Means are shown by a red cross, individual plant datapoints are shown by black open circles. Black lines connect the lowest and highest value in each genotype. Letters at the top of each plot indicate statistically different groups (p<0.05, one-way ANOVA and TukeyHSD).

### *ZmGLK1* and *ZmG2* transcript levels are reduced in *Zmscr1;Zmscr1h* mutants

Lower overall chlorophyll levels may reflect an underlying reduction in chloroplast size or number, in either M or BS cells of *Zmscr1;Zmscr1h* double mutants. In maize, genes encoding the Golden-2 like (GLK) transcription factors ZmGLK1 and ZmG2 are subfunctionalised in that *ZmGLK1* transcripts accumulate preferentially in M cells and *ZmG2* transcripts accumulate in BS cells (Rossini et al., 2001). Loss of ZmG2 function specifically perturbs BS chloroplast development (Hall et al., 1998; Langdale and Kidner, 1994). We have previously shown that *ZmSCR1* and *ZmSCR1h* transcripts accumulate most strongly in the ground meristem cells of leaf primordia that give rise to M cells in the mature leaf (Hughes et al., 2019). We therefore hypothesised that if ZmSCR1 and/or ZmSCR1h had a direct effect on chloroplast development, *ZmGLK1* but not *ZmG2* transcript levels would be reduced in *Zmscr1;Zmscr1h* mutant leaf primordia. Consistent with this hypothesis, qRT-PCR analysis revealed that *ZmGLK1* transcript levels were significantly reduced by on average 62% (*m2m1*) or 72% *(m2m2*) in double mutants (Fig 2A). Although there was some biological variation between individuals of both wild-type and mutant lines, this pattern was consistent and was replicated in two independent experiments. By contrast, levels of *ZmG2* transcripts were only significantly different from wild-type in the *m2m1* line, and even in this line the reduction was far less severe (reduced by only 43%) than seen with *ZmGLK1* (Fig 2A). Together these results suggested a direct or indirect role for ZmSCR1 and ZmSCR1h in the regulation of *ZmGLK1*, and thus possibly in the development of M cell chloroplasts.

**Figure 2.**
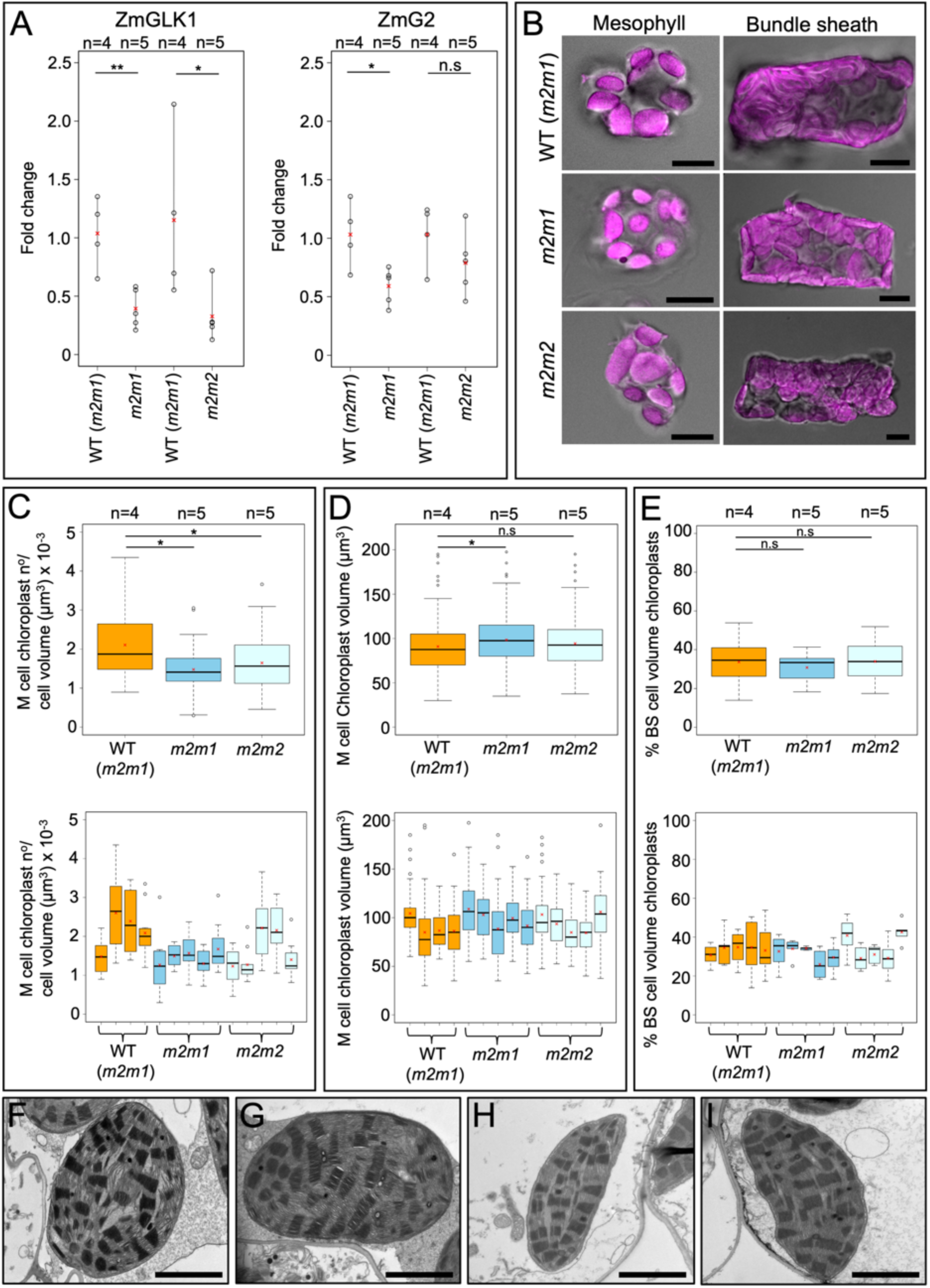
Chloroplast development is not perturbed in *Zmscr1;Zmscr1h* mutants. **A)** Relative transcript levels expressed as fold change of *ZmGLK1* (left) and *ZmG2* (right) transcripts in both *m2m1* and *m2m2* double mutants. Biological replicates (n=) are shown above each genotype. Means are shown by a red cross, individual plant datapoints are shown by black open circles. Black lines connect the lowest and highest value in each genotype. Statistical significance between WT (*m2m1*) and mutants was assessed using a one-way ANOVA: \**P*≤0.05; \*\**P*≤0.01; n.s.*P*≥0.05. **B)** Isolated mesophyll and bundle-sheath cells from each genotype. Chloroplast autofluorescence is pink. Scalebars are 10μm. **C)** Chloroplast number normalised by estimated cell volume in M cells. **D)** M chloroplast volume. **E)** % BS cell volume occupied by chloroplasts. In **C-E**, top plots are pooled results from all biological replicates, bottom plots are data from each plant assessed. Means are shown by a red cross, median by a horizontal black line. Statistical significance was assessed using a one-way ANOVA and TukeyHSD: \**P*≤0.05; n.s.*P*≥0.01. **F-I)** TEM images of M cell chloroplasts for both WT (*m2m1*) (F&G) and *m2m1* mutants (H&I). Scalebars are 2μm.

### Chloroplast development is largely unaltered in *Zmscr1;Zmscr1h* mutants

To assess whether reduced *ZmGLK1* transcript levels in *Zmscr1;Zmscr1h* double mutants led to perturbed chloroplast development, M cells were isolated from the leaves of both WT and *Zmscr1;Zmscr1h* mutants (Fig 2B), and both size and number of chloroplasts were quantified. M cell chloroplast numbers were counted in 7 to 10 cells from four WT (*m2m1*), five *m2m1* and five *m2m2* plants, and values were normalised by estimated cell volumes. Figure 2C shows that there was a statistically significant reduction in the number of chloroplasts per M cell in *Zmscr1;Zmscr1h* mutants, however, this reduction is extremely subtle and was not visually obvious (Fig 2B). Furthermore, when the data were viewed on a plant by plant basis it became clear that there was variation between plants of the same genotype, particularly in the *m2m2* mutant line where two of the five individuals showed no apparent average reduction in chloroplast number per cell (Fig 2C). Sampling more plants might reduce noise in this dataset, however, single-cell isolation, imaging and subsequent quantification on greater numbers than reported here would be prohibitively time-consuming. To assess whether individual M chloroplast size was altered in *Zmscr1;Zmscr1h* mutants, the volume of 10 chloroplasts per cell was measured in five distinct cells from four WT (*m2m1*), five *m2m1* and five *m2m2* plants. A subtle but statistically significant increase in average chloroplast volume was found in *m2m1* but not *m2m2* plants (Fig 2D), indicating that the slightly lower number of chloroplasts per cell in *m2m1* plants (Fig. 2C) is compensated for by an increase in the volume of individual chloroplasts. Finally, to determine whether M chloroplast ultrastructure was perturbed in the absence of changes in number or size, transmission electron microscopy was undertaken on mature leaves of both WT (*m2m1*) and *m2m1* mutant plants. M chloroplasts in WT (*m2m1*) (Fig 2F & 2G) and *m2m1* mutants (Fig 2H & 2I) were indistinguishable, with *m2m1* chloroplasts exhibiting normal thylakoid structure and grana. As such, the 60-70% reduction in *ZmGLK1* transcript levels in leaves of *Zmscr1;Zmscr1h* mutants does not impact on total chloroplast volume per M cell, or on chloroplast ultrastructure.

Although lower in magnitude and less consistent between the *m2m1* and *m2m2* mutant lines, the reduction in *ZmG2* transcript levels in *Zmscr1;Zmscr1h* leaves could feasibly perturb BS chloroplast development. Maize BS cells contain many tightly packed centrifugal chloroplasts such that counting or measuring individual chloroplasts in single cells is error-prone. As such, a thresholding method was used instead. Specifically, the total % of BS cell volume that contained chloroplasts was measured in 5 to 10 individual BS cells from five individual plants for each genotype. No significant difference was found between WT (*m2m1*) and *Zmscr1;Zmscr1h* mutants, with % chloroplast volume per cell being 30% to 35% in both (Fig 2E). These data indicate that the 43% (*m2m1*) reduction in *ZmG2* transcript levels in leaves of *Zmscr1;Zmscr1h* mutants is not sufficient to induce the changes in BS chloroplast development that are observed in loss of function *Zmg2* mutants. Collectively, these data reveal that lower chlorophyll levels in *Zmscr1;Zmscr1h* mutants are not associated with perturbed chloroplast biogenesis in M or BS cells, suggesting that the reduction may result from direct perturbations to the chlorophyll biosynthetic pathway, components of which are direct targets of GLK transcription factors in Arabidopsis (Waters et al., 2009).

### *Zmscr1;Zmscr1h* operate a normal C_4_ photosynthetic cycle with reduced capacity

Given that growth and chlorophyll levels are perturbed in *Zmscr1;Zmscr1h* mutants but chloroplast development is largely unaffected, we sought to determine whether photosynthetic capacity was compromised. As an initial screen, the maximal rate of carbon assimilation (A_max_) was measured under saturating CO_2_ conditions. Figure 3A shows that in both *Zmscr1;Zmscr1h* mutant lines, A_max_ was on average around 20 µmol m^-2^ s^-1^ as compared to around 30 µmol m^-2^ s^-1^ in wild-type. This reduction was consistent across all plants measured (n=6 for each genotype), across two independent experimental set-ups (Supplemental Figure 2). To test whether *Zmscr1;Zmscr1h* mutants also had reduced photosynthetic capacity under atmospheric conditions (400 μmol mol^-1^ CO_2_), A/C_i_ (CO_2_ assimilation, A, versus substomatal CO_2_ concentration, C_i_) curves were generated for at least three different plants from each genotype (Fig 3B). At 400 μmol mol^-1^, C_i_ levels were similar between genotypes, indicating that stomatal behaviour was not significantly altered (Fig 3B). However, photosynthetic rates were reduced by a similar magnitude to that seen at saturating CO_2_, with rates only slightly below the A_max_ rate in both wild-type and mutants (Fig 3B). This reduction in photosynthetic capacity likely underpins the perturbed growth phenotype in *Zmscr1;Zmscr1h* mutants.

**Figure 3.**
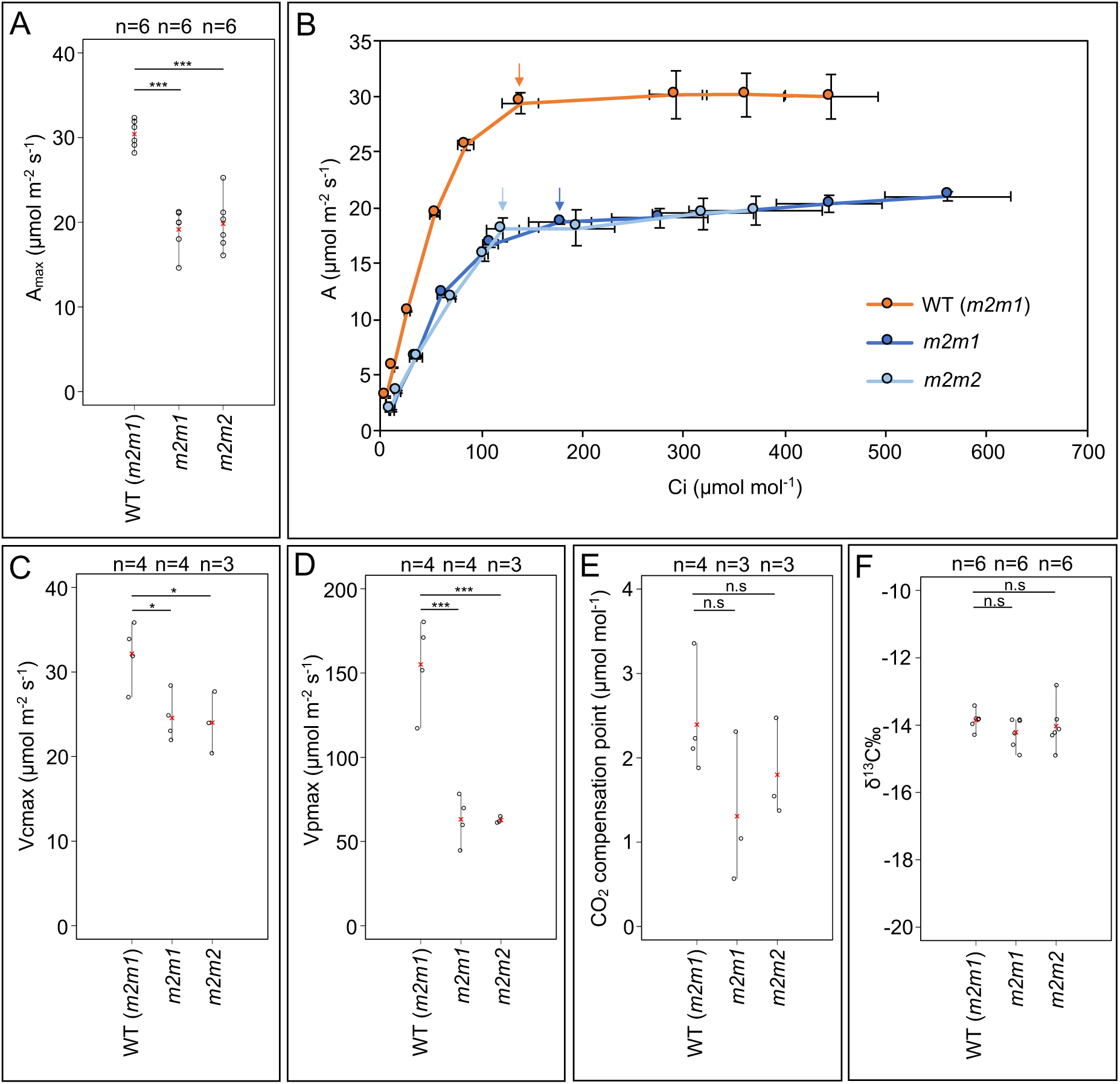
Photosynthesis is perturbed in *Zmscr1;Zmscr1h* mutants. **A)** A_max_ recorded at 800 μmol mol^-1^ CO_2_. **B)** A/Ci curves for each genotype. Each data point is the mean of at least three curves from different plants. Arrows within the plot indicate ambient measurements at 400 μmol mol^-1^. Error bars are standard errors of the mean. **C)** V_cmax_. **D)** V_pmax_. **E)** CO_2_ compensation point. **F)** δ^13^C ‰. In (A) and (C-F) biological replicates (n=) are shown above each genotype. Means are shown by a red cross, individual plant datapoints are shown by black open circles. Black lines connect the lowest and highest value in each genotype. Statistical significance between WT (*m2m1*) and mutants was assessed using a one-way ANOVA: \**P*≤0.05; \*\**P*≤0.01; \*\*\**P*≤0.001;n.s.*P*≥0.05.

In mature leaves of both *Zmscr1;Zmscr1h* mutant lines, the majority (>50% in both cases) of veins are separated by only one M cell rather than the normal two, skewing the 1:1 ratio of BS:M cells assumed to be critical for efficient C_4_ photosynthesis in wild-type maize and other Kranz exhibiting C_4_ species (Hughes et al., 2019). To determine whether the relative reduction in M cell volume leads to reduced PEPC-mediated carbon fixation capacity in *Zmscr1;Zmscr1h* mutants, we modelled the A/C_i_ curves using previously established C_4_ photosynthesis equations (von Caemmerer, 2000). These modelled assimilation curves enabled the estimation of two key parameters, the maximal rate of RuBisCO carboxylation (V_cmax_) and the maximal rate of PEPC carboxylation (V_pmax_). Figure 3C shows that V_cmax_ was reduced in both mutant lines, consistent with the lower maximal rates of photosynthesis observed. V_pmax_ estimation, which is based on the initial slope of the A/C_i_ curve, was also reduced in the mutants compared to the wild-type (Fig 3D). The estimated V_pmax_ in *Zmscr1;Zmscr1h* mutants was around a third of that found in wild-type (Fig 3D), indicating a significant reduction in PEPC capacity at a whole leaf level.

Given the reduction in PEPC capacity in *Zmscr1;Zmscr1h* mutants, alongside the perturbed Kranz patterning, it seemed plausible that some CO_2_ could be fixed directly by RuBisCO in the BS in a C_3_ cycle. To test this, we extrapolated A/C_i_ curves to estimate the CO_2_ compensation point in wild-type and *Zmscr1;Zmscr1h* plants. This analysis revealed no significant difference in the CO_2_ compensation point between wild-type and *Zmscr1;Zmscr1h* mutants (Fig 3E). We also carried out a δ^13^C ‰ analysis, which exploits the fact that PEPC and RuBisCO discriminate differently between ^12^C and ^13^C. Fixation by PEPC in C_4_ species results in δ^13^C ‰ values between −12 and −16, whereas fixation by RuBisCO in C_3_ species gives values between −20 and −37 (O’Leary, 1981). Notably, consistent with the CO_2_ compensation point estimations, δ^13^C ‰ values in *Zmscr1;Zmscr1h* mutants were the same as WT (Fig 3F). Collectively, our results indicate that *Zmscr1;Zmscr1h* plants operate a normal C_4_ photosynthetic cycle at significantly reduced capacity.

## Discussion

The coordination of growth and development are of fundamental importance throughout the plant lifecycle. SCARECROW is one of the best studied plant developmental regulators, with roles in patterning processes in both roots and shoots (Cui et al., 2014; Di Laurenzio et al., 1996; Wysocka-Diller et al., 2000). However, comparatively little attention has been paid to the perturbations in growth and physiology which accompany patterning defects in loss of function *scr* mutants. Here, we have demonstrated that reduced chlorophyll levels in maize *Zmscr1;Zmscr1h* double mutants (Figure 1) are not associated with changes in chloroplast development, despite a reduction in transcript levels of *ZmGLK1* and to a lesser extent *ZmG2* (Figure 2). We have also shown that *Zmscr1;Zmscr1h* mutants have reduced photosynthetic capacity both in terms of maximal overall rate and PEPC carboxylation rate, but that the plants operate a normal C_4_ cycle despite the associated Kranz patterning defects (Figure 3). Either directly or indirectly, SCARECROW therefore regulates photosynthetic capacity in maize.

There is some evidence to support a direct role for *SCARECROW* genes in regulating plant physiology. For example, the Arabidopsis *SCARECROW-LIKE (SCL)* GRAS proteins SCL6/22/27 directly regulate light-induced chlorophyll biosynthesis in a gibberellin-dependent manner (Ma et al., 2014), and protochlorophyllide oxidoreductase B, which catalyses one of the latter steps in chlorophyll biosynthesis, is a putative direct target of AtSCR itself (Cui et al., 2014; Garrone et al., 2015). AtSCR also directly modulates the sugar response (Cui et al., 2012) and genes encoding components of the light harvesting complex of photosystems I and II are putative targets (Cui et al., 2014). Most interestingly, AtSCR is proposed to bind to the promoter of *AtGLK2* but not *AtGLK1* (Cui et al., 2014). In light of these data, and given the functional conservation between maize and Arabidopsis SCARECROW (Lim et al., 2005), it is plausible that ZmSCR1/1h directly regulates expression of *ZmGLK1* and/or *ZmG2*.

In *Zmscr1;Zmscr1h* mutants, *ZmGLK1* transcript levels are reduced to a greater extent than *ZmG2* levels on a whole leaf basis. This observation is consistent with the suggestion that ZmSCR1 could directly activate *ZmGLK1* gene expression because *ZmSCR1* and *ZmSCR1h* are expressed predominantly in M cell precursors (Hughes et al., 2019). Some of the reduction in *ZmGLK1* levels can be attributed to the altered M cell patterning that results in >50% vascular bundles being separated by only one M cell rather than the normal two. However, this anatomical difference could only explain a 25% reduction in transcript levels at the whole leaf level whereas the observed reduction was 62% (*m2m1*) or 72% (*m2m2*). As such there is a ∼35-40% reduction in *ZmGLK1* transcript levels that is independent of any patterning defects, a similar reduction to that seen for *ZmG2* transcripts. Given that *ZmG2* is expressed preferentially in BS cells, the most parsimonious explanation for this observation is that loss of ZmSCR1/ZmSCR1h function has an indirect effect on *ZmGLK1/ZmG2* expression, although the possibility of direct but non-cell autonomous activation cannot be excluded. In either case, it can be concluded that *ZmGLK1* and *ZmG2* levels are high enough in *Zmscr1;Zmscr1h* mutants to facilitate normal chloroplast development.

Given the marked reduction in total chlorophyll in *Zmscr1;Zmscr1h* mutants (∼50% of wild-type), it is counter-intuitive that chloroplast development is apparently normal because intermediates in the chlorophyll biosynthetic pathway are known to participate in retrograde signalling pathways from the chloroplast to the nucleus (Larkin, 2016; Larkin et al., 2003; Mochizuki et al., 2001). It is possible that ZmSCR1 and ZmSCR1h affect chlorophyll biosynthesis at a point in the pathway downstream of the intermediates responsible for retrograde signalling and the suppression of chloroplast development. However, this is unlikely given the reduction in *ZmGLK1* and *ZmG2* transcript levels, because the Arabidopsis orthologs *AtGLK1* and *AtGLK2* directly activate chlorophyll biosynthesis genes throughout the pathway (Waters et al., 2009). An alternative possibility is that ZmSCR1 and ZmSCR1h play a direct or indirect role in the retrograde signalling pathway itself, such that loss of function prevents feedback from the chloroplast to the nucleus. In either case, our results indicate that at the level of chlorophyll reduction observed, it is possible to uncouple chlorophyll biosynthesis and chloroplast development.

Although both chloroplast development and the C_4_ photosynthetic signature appear normal in *Zmscr1;Zmscr1h* mutants, lower chlorophyll levels are accompanied by reduced maximum photosynthetic capacity (A_max_), RuBisCO carboxylation rate (V_cmax_) and PEPC-carboxylation rate (V_pmax_). Reduced V_pmax_ may be a reflection of the relative reduction in total M cell volume that results from the known patterning defect. However, the reduction in V_cmax_ is unlikely to result from anatomical changes because BS cells, and the chloroplasts within them, are unaltered (Hughes et al., 2019). It is more likely that the observed reduction in overall photosynthetic capacity results from a reduction in the amount of light that can be harvested to power photosynthesis as a consequence of the lower chlorophyll levels. Collectively, our results demonstrate that in addition to well-established patterning roles, SCR helps to establish and/or maintain photosynthetic capacity in maize.

## Materials and methods

### Plant material and growth conditions

Both *Zmscr1-m2;Zmscr1h-m1* (*m2m1*) and *Zmscr1-m2;Zmscr1h-m2* (*m2m2*) double mutants were generated and described previously (Hughes et al., 2019). As segregating wild-type lines for both were indistinguishable, the WT (*m2m1*) line was chosen for comparison of both lines in this study. Plants were germinated and grown in a greenhouse in Oxford, UK with a 16 h/8 h light/dark cycle, with supplemental light provided when natural light dropped below 120 μmol photon m^−2^ s^−1^. Day temperature was 28°C and night temperature was 20°C. Seeds were germinated in vermiculite and transferred after 9-11 days to 12 cm pots containing 3:1 mixture of John Innes No. 3 Compost (J. Arthur Bower) and medium vermiculite (Sinclair Pro). After 24 days, plants were re-potted in 30 cm pots using the same soil mixture with the addition of Osmocote Exact Standard 3-4M (ICL) slow release fertiliser. Plants were arranged randomly for the duration of growth to eliminate greenhouse location effects.

### Chlorophyll extraction and measurements

A 0.25 cm^2^ leaf disk was taken from ¾ along the proximal-distal axis of leaf 4 23 days after planting, avoiding the midrib. The leaf disk was frozen in liquid nitrogen, homogenised and resuspended in 2 ml of ice-cold 80% (v/v) acetone before centrifugation (17900 x *g*) for 5 min at 4°C. The supernatant was then removed and kept on ice in the dark prior to measurements. 1 ml of supernatant was transferred to a 1.5 ml quartz cuvette and absorbance recorded at 646.6nm, 663.6nm and 750nm. Total chlorophyll, chlorophyll a and chlorophyll b contents were calculated on a leaf area basis as described previously (Porra et al., 1989).

### Quantitative RT-PCR analysis

Two 0.25 cm^2^ leaf disks per plant were harvested from developmentally equivalent immature leaves emerging from the whorl 31 days after planting, and frozen in liquid nitrogen. Total RNA was extracted using an RNeasy Plant Mini Kit following the manufacturer’s instructions (Qiagen). RNA was then treated with DNaseI (Invitrogen), and 1 µg of DNase-treated RNA was used as template for cDNA synthesis using a Maxima First Strand cDNA synthesis kit (Thermo Scientific). cDNA quality was assessed by RT-PCR using primers amplifying a product with distinct sizes in cDNA and genomic DNA extracts.

Quantitative RT-PCR (qRT-PCR) experiments were undertaken using SYBR Green. Cycle conditions were 95°C for 10 minutes, then 40 cycles of 95°C for 15 seconds and 60°C for 1 minute. Melt curves were generated between 60°C and 95°C to confirm a single product was amplified (Supplemental Figure 1). Primers were designed for the CDS of both *ZmGLK1* (GRMZM2G026833) and *ZmG2* (GRMZM2G087804) using Primer3Plus (Untergasser et al., 2012), amplifying regions ∼100bp in length (Supplemental Figure 1). Primers were first tested by RT-PCR using Gotaq (Promega) with cycle conditions 95°C for 5 min, 35 cycles of 95°C for 30 s, 57°C for 30 s and 72°C for 30 s, and 72°C for 5 min, to confirm correct amplicon size (Fig S1B). qRT-PCR amplification efficiencies were calculated using 1/4, 1/8, 1/16, 1/32 and 1/64 wild-type cDNA dilutions. Experiments were run with primers for two housekeeping genes, *ZmCYP* and *ZmEF1α*, that have been previously validated as suitable controls in maize leaf experiments (Lin et al., 2014). Three technical replicates were undertaken for each sample and confirmed to have Ct values with range <0.5. All relevant comparisons were run on the same plate alongside water controls, and as such the WT (*m2m1*) samples were repeated alongside the *m2m2* samples. Ct values were calculated using the real-time PCR miner algorithm (Zhao and Fernald, 2005), and fold-change values were calculated using the 2^-ΔΔCT^ method (Livak and Schmittgen, 2001). Each wild-type sample was compared individually to the average wild-type values in order to indicate the spread of the wild-type data. Mutant samples were then compared to the same wild-type average.

### Single cell isolation and chloroplast quantification

Strips of leaf ∼1 cm long and ∼1-2 mm wide were cut from the mid-point along the proximal-distal axis of fully expanded leaf 5, 33 days after planting. Leaf strips were harvested and fixed in 0.5% glutaraldehyde in PBS (0.15 M NaCl, 10 mM phosphate buffer pH 6.9), for 1 hour in the dark at room temperature as previously described (Wang et al., 2017). Samples were then stored in Na_2_EDTA buffer (0.2 M disodium EDTA, pH 9) at 4°C until digestion. For single cell isolation, samples were incubated in Na_2_EDTA buffer for 3 hours at 55°C, before being rinsed in digestion buffer (0.15 M sodium hydrogen phosphate, 0.04 M citric acid, pH 5.3) twice for 15 minutes. Samples were then incubated at 45°C in 2.5% pectinase in digestion buffer for 1 hour, before rinsing twice in digestion buffer. All digested samples were stored at 4°C and imaged within a week.

For imaging, digested leaf strips were placed on a glass slide in a drop of 1:1 Na_2_EDTA buffer: glycerol. The bottom of a 1.5 ml eppendorf tube was used to apply pressure to the leaf strip in order to separate the cells. For the isolation of single M cells, light pressure was applied by tapping the tube bottom on the edges of the leaf strip, whereas more force was applied for the isolation of single BS cells. Images were acquired using a Leica SP5 laser scanning confocal microscope and Leica Application Suite (LAS) software. Chlorophyll was excited at 561 nm and emission collected between 650 nm and 750 nm. Transmitted laser light provided pseudo bright field images. Cells were imaged in each focal plane at 500 nm step size such that a z-stack comprising the entire cell was generated.

Image quantification was undertaken using Fiji (https://imagej.net/Fiji). M cell volumes were calculated by assuming a cuboid shape, whereas BS cells were treated as cylinders. M chloroplast volumes were calculated using the volumest plugin in Fiji. M cell chloroplast counts were undertaken blind such that the genotypes were unknown until after quantification was complete. Total BS chloroplast volumes were calculated by thresholding the image using the Otsu method. Minimum and maximum threshold parameters were automatically calculated, although in some cases manual adjustment was necessary to ensure all chloroplasts were included. The area of threshold pixels was then counted in each z-stack layer before being combined for an estimated total chloroplast volume per cell.

### Transmission electron microscopy

Leaf 4 was sampled with a 2 mm leaf punch at the mid-point along the proximal-distal axis 29 days after planting. Leaf disks were cut in half and then fixed in a microwave fixation unit (Leica EM AMW) with 2.5% glutaraldehyde 4% paraformaldehyde in 0.1 M sodium cacodylate buffer, pH 6.9, for 4 × 2 mins at 37°C, alternating 20 W and 0 W microwave power (continuous), before final incubations for 2 min with 30 W (pulse), 2 min with 0 W power and 4 min with 20 W (pulse), all at 37°C. Samples were then placed in fresh fixative at 4°C for 3 days. Unless stated all subsequent microwave power settings were continuous rather than pulsed.

Samples were then further processed by microwave fixation by incubating for 4 × 1 min (37°C, 20 W) in 0.1 M sodium cacodylate buffer, pH 6.9, then for 12 min (37°C, 20 W) in 2% osmium tetroxide and 1.5% potassium ferricyanide in the same cacodylate buffer. Samples were then washed in dH_2_O for 9 × 1 min (37°C, 15 W), before being incubated in 2% uranyl acetate for 5 min (37°C, 20 W pulse), then 2 min at (20°C, 0 W) and 2 min at (37°C, 15 W) and washed 3 × 1 min (37°C, 15 W pulsed) in dH_2_O. Following this, samples were dehydrated through 10%, 20%, 30%, 40% (all 20°C, 1 min, 10 W, v/v), 50%, 60%, 70% (x2), 80% and 90% (37°C, 1.5 mins, 25 W, v/v) ethanol. To complete the dehydration, samples were incubated in 100% ethanol for 2 × 2 mins, 1 × 1 hour (all 37°C, 25 W) and 2 × 2 min (37°C, 25 W), 1 × 1 min (37°C, 10 W), 2 × 2 min (37°C, 25 W) and 1 × 1 min (37°C, 10 W) 100% acetone (v/v). Samples were resin infiltrated with TAAB Hard Plus resin for 3 min each in 25% (37°C, 10 W), 50% (37°C, 10 W) and 75% (45°C, 12 W) dilutions made up with acetone (v/v). Samples were then incubated in 100% TAAB Hard Plus resin for 3 min (45°C, 12 W), 2 × 10 min (45°C, 12 W), 2 × 15 min (45°C, 12 W) and 30 min (50°C, 12 W). Finally, samples were removed from the microwave fixation unit and polymerised in moulds with fresh resin for 36 hours at 65°C. For sectioning, blocks were trimmed and 90nm sections cut using a diamond knife. Sections were post-stained with 2% lead citrate for 5 minutes before rinsing with degassed dH_2_O four times. Sections were then dried and imaged using a FEI Tecnai 12 transmission electron microscope operated at 120kV with a Gatan OneView camera.

### Photosynthesis measurements

Steady state A_max_ measurements and A/Ci curves were generated using an infrared gas analyser (Li-6800, LI-COR, USA) with a 2 cm^2^ leaf chamber fluorometer head. A_max_ measurements were undertaken in the greenhouse on fully expanded leaf 8, 49 days after germination, at 800 μmol mol^-1^ CO2, 1800 μmol photons m^−2^ s^−1^, 60% humidity and 26°C leaf temperature. Leaves were clamped at the midpoint along the proximal-distal axis, adjacent to but not including the midrib. Leaves were allowed to acclimate for around 30 mins until photosynthetic rate had stabilised, at which point A_max_ was recorded. Measurements were taken throughout the day between 10:00 and 18:00, with genotypes randomly sampled to avoid any circadian influences. A/C_i_ curves were generated using fully expanded leaf 6 between 29 and 32 days after planting. Plants were moved into the lab where leaves were clamped in the same way as for A_max_ measurements and allowed to acclimate for at least 30 minutes at 400 μmol mol^-1^ CO2, 1800 μmol photons m^−2^ s^−1^, 60% humidity and leaf temperature 26°C. A/C_i_ curves were generated by measurement of photosynthetic rate at 400, 300, 200, 100, 50, 25, 400 (recovery, omitted from curve plotting), 600, 800 and 1000 μmol mol^-1^ CO_2_. At each new CO_2_ concentration, measurements were only taken at steady state, normally achieved between 2 and 6 mins after the desired CO_2_ concentration was achieved.

Individual A/C_i_ curves were modelled using the equations described previously for C_4_ photosynthesis (von Caemmerer, 2000). The most recent kinetic constants described for *Setaria viridis* (also an NADP-ME C_4_ monocot) were used in order to fit the model and estimate both V_cmax_ and V_pmax_ (Boyd et al., 2015). Modelled data agreed closely with experimental data. Compensation points were estimated by fitting a simple polynomial trendline to the first 3-4 points of the modelled data, and solving this to estimate the x-axis intercept and thus the CO_2_ compensation point.

### δ^13^C‰ analysis

A small section of leaf tissue was sampled from leaf 10, 58 days after planting. This tissue was then freeze dried for 1 week. Once dry, tissue was ground to a powder by milling in a TissueLyser II (Qiagen) for 1 minute at 28 Hz. From this powder, 1 mg of tissue was weighed and placed in a tightly sealed tin capsule. ^13^C/^12^C ratios were measured using a Sercon 20/22 isotope-ratio mass spectrometry system coupled to a Sercon GSL elemental analyser. Samples were run alongside collagen standards which are traceable back to the VPDB standard. Known alanine controls were run alongside samples to ensure accuracy. δ^13^C ‰ was calculated as:

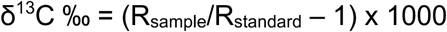

where R is the ratio of ^13^C mass to ^12^C mass.

## Acknowledgements

We thank Errin Johnson for help with the TEM analysis; Peter Ditchfield for δ13C ‰ analysis; Niloufer Irani for help with confocal imaging; Daniela Vlad, Chiara Perico, Sovanna Tan, Julia Lambret-Frotte and Roxaana Clayton for discussion throughout the experimental work and during manuscript preparation; Susanne von Caemmerer for help with A/C_i_ analysis; Bob Furbank for discussion of the manuscript; Phil Becraft, Hao Wu and Erik Vollbrecht for help crossing and maintaining mutant lines; and Julie Bull and Lizzie Jamison for technical support.

This work was funded by the Bill and Melinda Gates Foundation C_4_ Rice grant awarded to the University of Oxford (2015-2019; OPP1129902) and by a BBSRC sLoLa grant (BB/P003117/1).

## Author contributions

T.E.H and J.A.L conceived of the study and designed the experiments. T.E.H performed the experiments and analysed the data. T.E.H and J.A.L wrote the manuscript.

## Data statement

All relevant data is included in the manuscript and supporting information. Access to raw data is available on request.

---

**Supplemental Figure 1.**
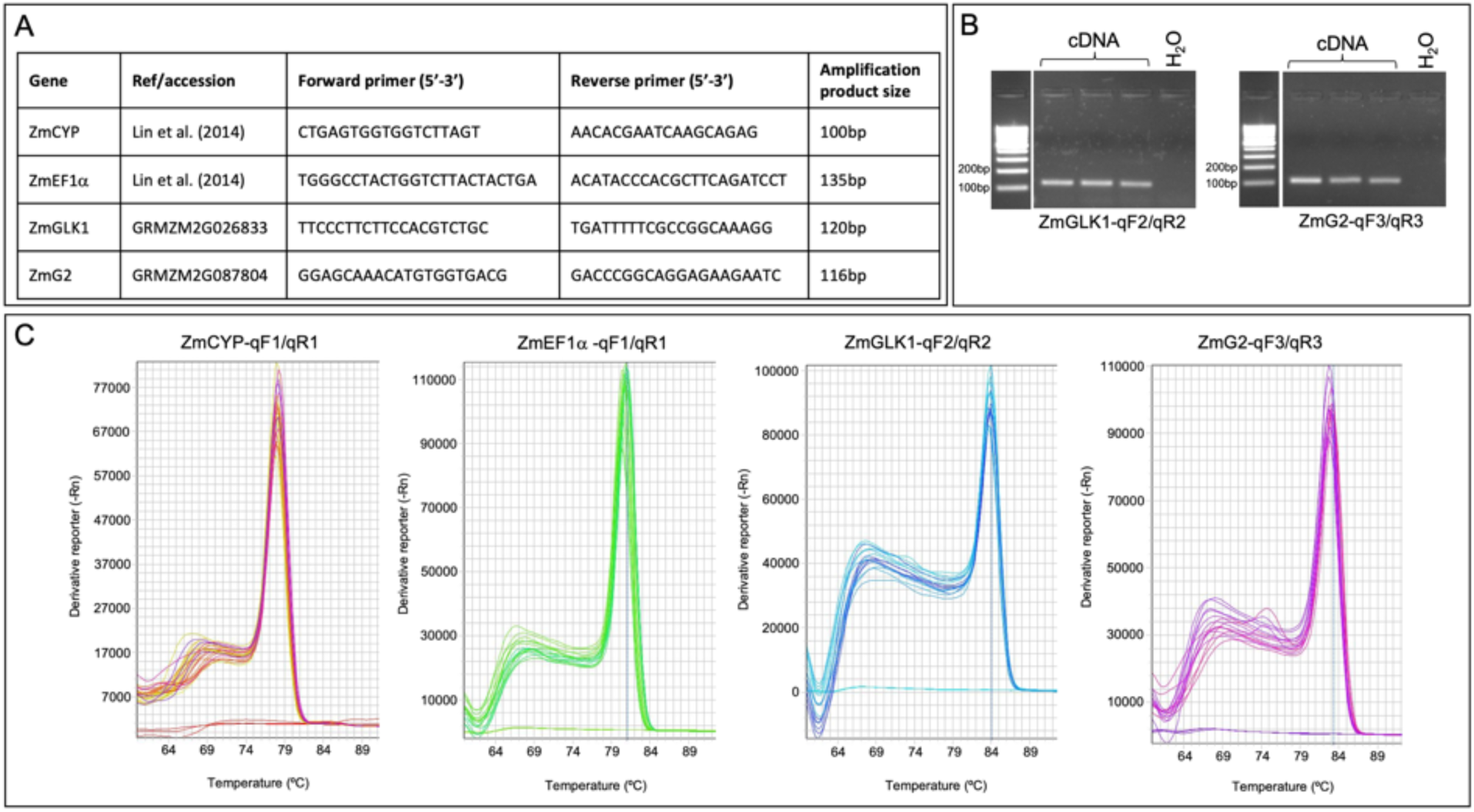
qRT-PCR validation. **A)** qRT-PCR primer design. **B)** *ZmGLK1* and *ZmG2* qRT-PCR primer testing by RT-PCR. Three cDNA samples from different plants are displayed for each primer pair. **C)** Melt curves of qRT-PCR amplification products for all primer pairs. Flat lines at the bottom are water controls.

**Supplemental Figure 2.**
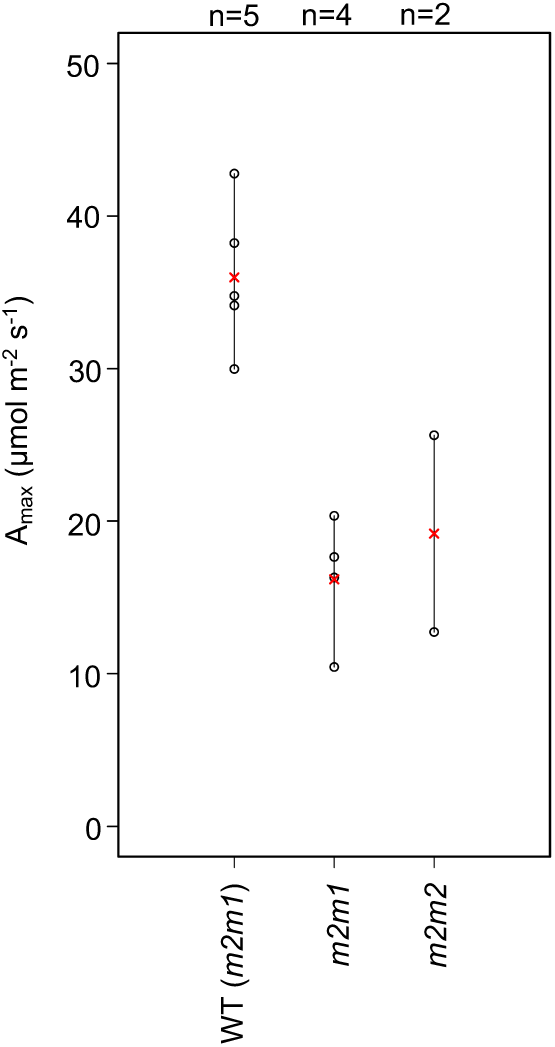
Independent A_max_ validation. Measurements were undertaken 35-38 days after planting on leaf 6.

